# A simple silicone elastomer colonisation model highlights complexities of *Candida albicans* and *Staphylococcus aureus* interactions in biofilm formation

**DOI:** 10.1101/2024.12.18.629256

**Authors:** Gail McConnell, Liam M. Rooney, Mairi E. Sandison, Paul A. Hoskisson, Katherine J. Baxter

## Abstract

Healthcare-associated infections (HAIs) significantly contribute to the burden of antimicrobial resistance (AMR). A major factor in HAIs is the colonisation of indwelling medical devices by biofilm-forming opportunistic pathogens such as *Candida albicans* and *Staphylococcus aureus*. These organisms frequently co-infect, resulting in synergistic interactions with enhanced virulence and resistance to treatment. *C. albicans* and *S. aureus* readily form dual-species biofilms on silicone elastomers, a commonly used medical device material, yet the colonisation phenotypes of these organisms on such surfaces remains poorly understood.

We developed a simple, optically tractable model to mimic the colonisation of indwelling medical devices to investigate *C. albicans* and *S. aureus* biofilm formation. The system utilises discs of a silicone elastomer embedded in agar, reflecting device-associated conditions and enabling high-resolution imaging of biofilms formed by *C. albicans* and *S. aureus* co-culture. Initial results using the silicone elastomer colonisation model reveal robust biofilm formation. These biofilms exhibited morphological differences between dual species biofilms formed by *S. aureus* co-cultures with either yeast- or hyphal-form *C. albicans*, indicating the impact of differing *C. albicans* cell morphotypes in biofilm-associated medical device colonization on silicone elastomers. Quantification of biofilm formation by crystal violet staining provided further validation of the system. These findings underscore the importance of developing tools for biofilm study which more closely resemble the infectious microenvironment, with our work detailing such a system which can be employed in further study to improve strategies against device-related HAIs.

## Introduction

Healthcare acquired infections (HAI) are recognised by the World Health Organisation as key contributors to the burden of antimicrobial resistance (AMR) (1) and result in longer hospital stays, higher levels of morbidity and increased economic cost (2,3). European HAI prevalence is 6.5% (4) and occurs at an incidence rate of 5-10% worldwide (5), highlighting the global burden of such infections. Treatment regimens are limited due to their association with multidrug-resistant microorganisms (3,6,7), and HAIs lead to more than 90,000 deaths in the EU per year alone (7).

A major risk factor for HAIs is the presence of invasive and indwelling medical devices (7–9) which act as a point of entry for microorganisms into the body and a route for systemic infection (10–12). Biofilms readily form on such indwelling medical devices including central venous catheters, indwelling stents, prosthetic heart valves and urinary catheters (10,13,14). They act as a source of chronic infection through dispersal of cells during the biofilm lifecycle (15) and can result in systemic infection including the formation of infectious metastases at distal sites (16,17). The only way to prevent resurgent infection is biofilm removal, normally removal of the device or implant at great risk to the patient (18).

The fungus *Candida albicans* and the bacterium *Staphylococcus aureus* are two opportunistic pathogens of the skin microbiome which regularly co-infect those with underlying health conditions (19,20). Intricate physical and metabolic interactions between *C. albicans* and *S. aureus* contribute to their synergy in infection (21–25) resulting in disease with greater recalcitrance to treatment than either organism alone (24,25) *C. albicans* and *S. aureus* are colonisers of silicone elastomers used as coatings or base materials for catheters, drainage tubing and other implantable medical devices (11,12,26), with *C. albicans* hyphae known to penetrate into silicone elastomers (27). To date, the structure of *C. albicans*/*S. aureus* dual-species biofilms on these materials remains to be fully investigated, leaving a critical gap in our knowledge around the behaviour of this important synergistic pairing in biofilm-associated medical device infections.

To improve our understanding of device colonisation by *C. albicans* and *S. aureus*, this study aimed to develop an optically transparent, low autofluorescence silicone elastomer microbial colonisation model to visualise biofilm formation by *C. albicans* and *S. aureus* co-culture. Model development required three criteria: namely the utilisation of inexpensive materials, the provision of conditions reflecting that of an indwelling medical device, and the ability to be coupled to an imaging system to provide high-resolution visualisation of biofilm. By applying these criteria, a simple silicone elastomer microbial colonisation model was developed using a 6-welled tissue culture plate in which discs of the widely used elastomer poly(dimethylsiloxane) (PDMS) were embedded in agar to provide a solid, nutritionally available substrate to represent incorporation of an indwelling device into tissue.

To identify the suitability of the silicone elastomer microbial colonisation model (from this point on referred to as the SIMCO model) for biofilm imaging, trials were performed on biofilms formed by *C. albicans* and *S. aureus* co-cultures using the Mesolens, a bespoke microscope system coupling low magnification with a high numerical aperture (28). This unique combination allows subcellular resolution across a maximum volume of capture of 6 mm x 6 mm x 3 mm (29) and enables high-resolution visualisations of whole mature colony biofilms. Recent applications of the Mesolens in microbiology have provided novel insight into biofilm macrostructures and biofilm community responses to nutrient availability and growth substrates (30–33). In terms of multispecies biofilms, Mesolens studies have revealed the emergence of macrostructures within *C. albicans* and *S. aureus* dual-species biofilms over time, and highlighted the impact of *C. albicans* cell morphotypes on biofilm architecture (34).

Using the above approach, images of robust biofilms formed by co-cultures on PDMS discs were captured using the Mesolens. Additional assessment of biofilm formation on the indwelling medical device mimic performed with crystal violet, a common staining method for biofilm quantification on surfaces (35,36), also supports its suitability for biofilm analyses. Critically, both these approaches revealed differences in biofilm formation between dual-species biofilms containing either predominantly yeast-form *C. albicans*, or hyphal-form *C. albicans*. As hyphal-form *C. albicans* is key in invasive disease, *S. aureus* interactions and medical device penetration (27,37,38), the ability to observe differences between yeast-form and hyphal-form *C. albicans* cell morphotypes in biofilm formation is paramount for development for novel approaches to medical device-related infection. The work presented here indicates the functionality and suitability of the SIMCOL model for imaging analysis of *C. albicans/S. aureus* biofilm formation, and details initial findings using the system.

## Methods

### Strains and growth conditions

Glycerol stocks of constitutively expressing fluorescent derivatives of *C. albicans* (pACT-1-GFP) (39) and *S. aureus* (N315 mCherry) (34) were respectively cultured on yeast extract peptone dextrose (YPD) and lysogeny broth (LB) agar (both Merck Life Science, UK). Generation of seed cultures for dual species co-cultures containing predominantly yeast-form *C. albicans* or hyphal-form *C. albicans* are described in detail in Baxter *et al* (2024) (34). Briefly, single colonies of either *C. albicans* or *S. aureus* were inoculated as monocultures in 5ml of lysogeny broth (Merck Life Science, UK). Cultures destined for hyphal-form co-culture were incubated at 37°C/250 rpm for 16 hours. Prior to overnight incubation of hyphal-destined cultures, aliquots of *C. albicans* and *S. aureus* monoculture were removed and diluted in a further 5 ml of lysogeny broth (500-fold for *C. albicans* and 1000-fold for *S. aureus* cultures) and incubated under the same conditions to generate cultures destined for yeast-form co-culture.

### Selection of components for an optically tractable silicone elastomer colonisation model

In terms of a suitable tissue mimic, previous work with LB supplemented with 0.2% glucose as a growth medium in the study of *C. albicans* and *S. aureus* dual-species biofilm formation (34) and its minimal autofluorescence under Mesolens imaging conditions (30,31) warranted its selection as a suitable tissue equivalent for our purposes. With regards to growth vessel needs, assessment criteria required a vessel with sufficient capacity for inclusion of both agar and culture volumes which would also allow ease of PDMS removal after culture incubation. Trials of both 12-welled and 6-welled tissue culture plates to assess volume capacity and PDMS removal indicated the greater suitability of 6-welled tissue culture plates. Finally, to identify differences in biofilm formation in the presence and absence of a tissue mimic, a PDMS-only control lacking LB agar was used in parallel. Figure 1 illustrates the design of the SIMCOL model.

**Figure 1:**
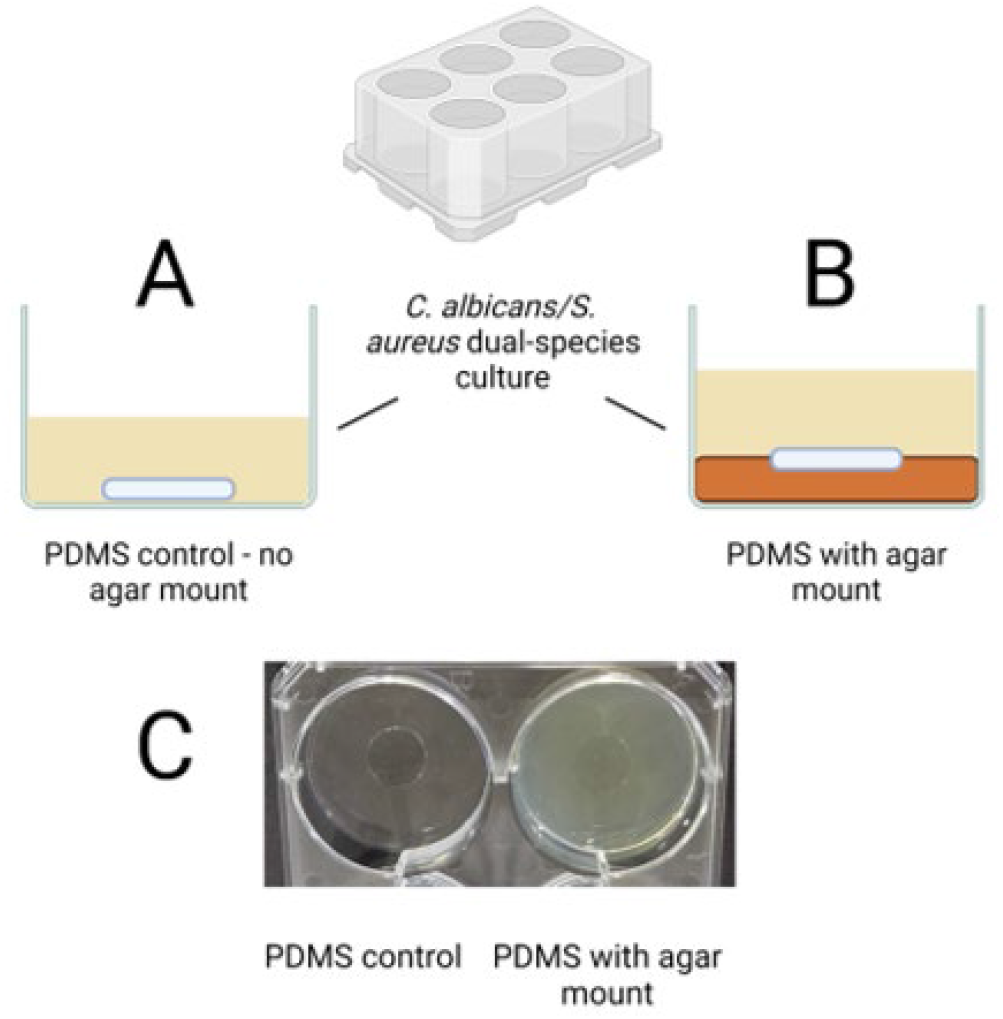
Silicone elastomer microbial colonisation (SIMCOL) model. **A:** Control PDMS discs were placed on the bottom of a well without agar. **B:** SIMCOL PDMS discs were partially embedded in 2 mL of molten agar at the bottom of a well of a 6-welled plate and left to solidify. **C:** Image of model set-up. After inoculation of 3 mL of culture, 6 welled plates were incubated at 37°C for 48 hours to allow biofilm formation. Figure created using BioRender.

PDMS discs were added to the model as shown in in Figure 1, subsequently removed after 48 hours and imaged using the parameters for GFP and mCherry widefield epifluorescence mesoscopy as described below. No significant background fluorescence was observed.

### Generation of PDMS discs

To produce PDMS discs, SYLGARD™ 184 Silicone Elastomer Base and curing agent (Dow Chemical company) were mixed at a ratio of 10:1 and poured onto a polished silicon wafer, which had been coated with a fluorosilane mould release layer (vapour deposition of trichloro(1H,1H,2H,2H-perfluorooctyl)silane, Merck Life Science UK), to give a final film thickness averaging 0.8 mm. PDMS was degassed in a vacuum desiccator and cured overnight at 50°C or for 48 hours at room temperature. After curing, the PDMS was removed from the silicon wafer and 13 mm diameter discs were cut from the PDMS sheet using a punch.

### Silicone elastomer microbial colonisation (SIMCOL) model setup

Prior to use, PDMS discs were immersed in 70% ethanol for 10 minutes, blotted on UV treated absorbent tissue and left to air dry in a microbiological safety cabinet. After drying, discs were placed on the upturned lid of a sterile 6-welled plate (ThermoScientific, UK) and irradiated with UV for 10 minutes on both sides. PDMS only control discs were placed in the bottom of one of the wells of a sterile 6-welled plate. For wells containing the indwelling medical device mimic, 2 ml of molten LB agar supplemented with 0.2% glucose was added to the bottom of another well of the same 6-welled plate, and a sterile PDMS disc immediately immersed. For generation of co-cultures, overnight cultures of *C. albicans* and *S. aureus* (described above) were diluted to an OD600 of 0.1 and 0.04, respectively, and cultures mixed at a ratio of 1:1 (v/v) to produce co-culture. A co-culture volume of 3 ml was then added to wells containing either the PDMS control or the indwelling medical device well. The 6-welled tissue culture lid was sealed with parafilm (Bemis, USA) and incubated at 37°C for 48 hours at 80rpm.

### Development of workflow for biofilm analysis

Previous studies of biofilm formation on surfaces detail the need for washing to remove nonadherent cells prior to investigation (40–42). Several different wash steps were trialled before optimising a regime which removed excess unattached cells without removing attached biofilms. Figure 2 demonstrates the optimal wash procedure devised for this investigation, and details the workflow undertaken before progressing to either (A) crystal violet staining or (B) widefield epifluorescence mesoscopic imaging.

**Figure 2:**
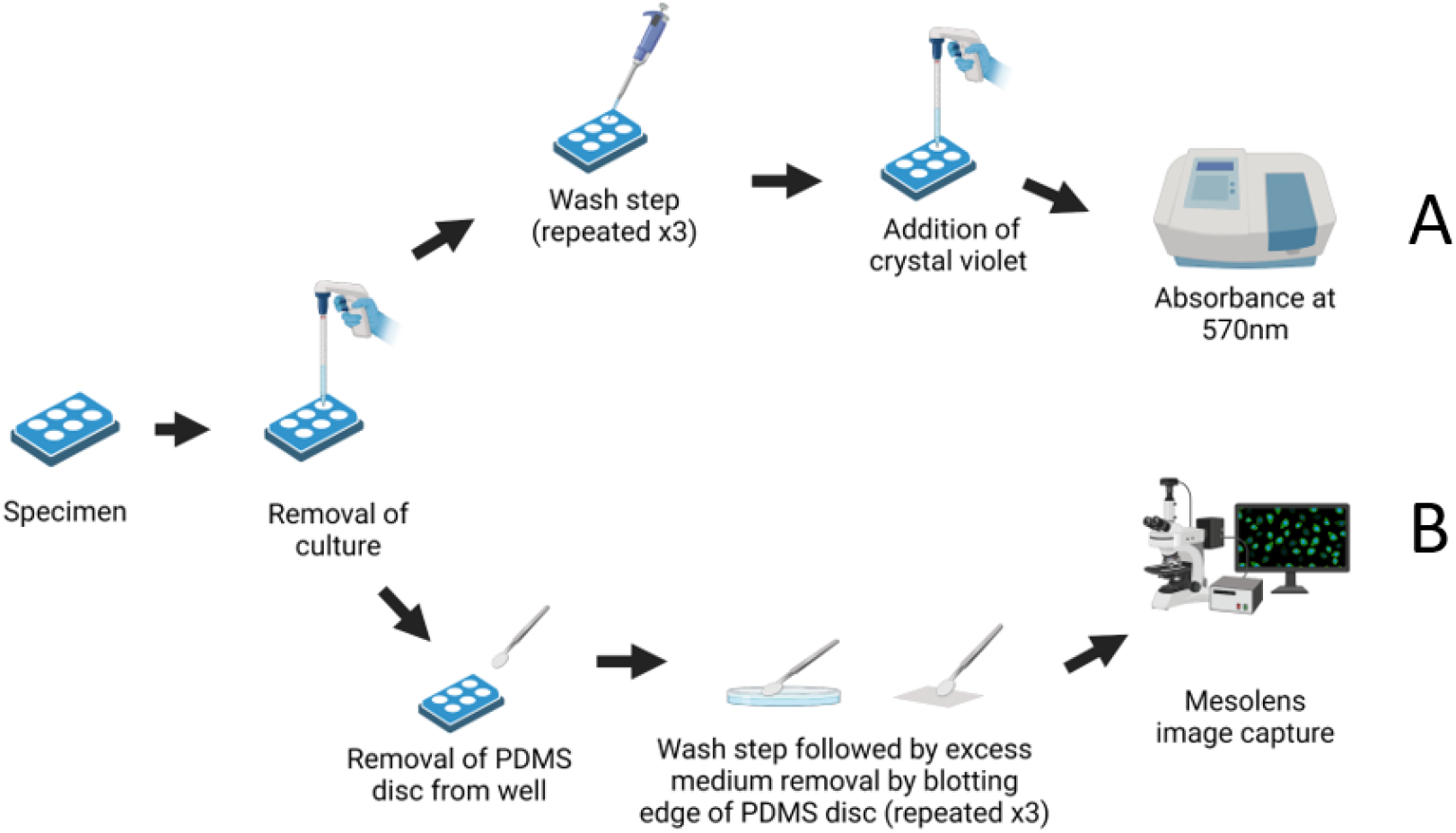
Workflow for assessment of biofilm formation on PDMS discs. Both crystal violet staining and mesoscopy involve washing PDMS discs in fresh medium prior to assay-specific downstream processing for (**A**) crystal violet staining of biofilms and (**B**) widefield epifluorescence mesoscopy. Figure created using BioRender.

### The silicone elastomer colonisation model as a system for following biofilm growth

Dual-species cultures containing either yeast-form or hyphal-form *C. albicans* were inoculated into the 6-welled plate containing the indwelling medical device mimic system and incubated for 48 hours at 37°C prior to analysis. Biofilm deposition was assessed by widefield epifluorescence mesoscopic imaging and matrix production investigated by crystal violet assay.

### Preparation of specimens for mesoscopy

After 48 hours incubation, the *C. albicans/S*.*aureus* co-culture was removed from the 6-welled tissue culture plate, and PDMS discs for mesoscopic imaging were excised from the 6-welled tissue culture plate. Excess culture was removed by gently placing the edge of the PDMS disc against an absorbent paper towel. PDMS discs were washed by careful immersion into phosphate buffered saline (PBS) (Oxoid, UK) at pH 7.3, and unattached cells were removed by gentle agitation. After washing, excess liquid was again removed from PDMS discs by gently blotting the edge of the disc against a paper towel. This wash process was repeated three times before discs were placed on a glass microscope slide (AGL4380-1 Agar Scientific, UK). A small amount of water was added to the surface of the PDMS discs before the addition of a glass coverslip (70×70 mm, type 1.5 (017999098; Marienfield, Lauda-Koenigshofen, Germany) before imaging.

### Widefield epifluorescence mesoscopy

To visualise *C. albicans* pACT-1 GFP and *S. aureus* N315 mCherry deposition on the PDMS surface, fluorescent proteins were excited using a CoolLED pE-4000 light emitting diode (LED) engine (CoolLED, UK) at excitation/emission wavelengths of 490 nm/525 nm ± 20 nm for GFP and 585 nm/635 nm ± 20 nm for mCherry. High resolution images were captured by a chip-shifting camera sensor, as detailed in Baxter *et al* (2024) (34), with the correction collars of the Mesolens set for water immersion. Z-stacks of biofilms were captured by moving the specimen axially in 5 µm increments using a computer-controlled specimen stage (Optiscan III, Prior Scientific,UK).

### Crystal violet staining

Crystal violet staining has been extensively used for the investigation and quantification of biofilm formation (35,36,41,43) as it binds negatively charged molecules of the biofilm matrix and is quantifiable by spectrophotometry at wavelengths of 550-570 nm. To quantify biofilm matrix deposition on the surface of PDMS, crystal violet staining using a method adapted from Zainal *et al* (2019) (43) was performed. After 48 hours of incubation, co-culture was discarded and PDMS discs were removed and placed in a fresh 6-welled plate. 3 mL of 0.1% (v/v) crystal violet solution (Pro-Lab, UK) was added to each well and incubated for 15 minutes. Crystal violet was removed and discarded, and PDMS discs left to air dry for 15 minutes, after which 95% (v/v) ethanol was subsequently added for a further 15-minute incubation to extract the bound dye. Three technical replicates of 200 µl for each biological replicate were added to a 96-well plate, and absorbance was quantified at 570 nm in a SPECTRAMAX 190 spectrophotometer (Molecular Devices, USA).

### Image analysis

Z stacks of hyphal-form and yeast-form biofilms on PDMS with and without agar were captured as described above and converted to maximum intensity projections using the open-source FIJI image processing software (ImageJ, version 1.54f (44)). Total Biofilm coverage on each PDMS surface was calculated as a percentage of field of view (FOV). To indirectly quantify and compare total biomass of *C. albicans* and *S. aureus* cells on the PDMS surface between growth conditions, the corrected fluorescence intensities of images of *C. albicans* and *S. aureus* were calculated as described previously in Baxter *et al* (2024) (34), and total biofilm cell mass on the PDMS surface derived from the addition of the corrected fluorescence intensities values obtained for both excitation wavelengths.

## Results

### Biofilm formation of dual species cultures on PDMS varies by both C. albicans cell morphotype, and the presence of agar

Widefield epifluorescence mesoscopy of biofilms formed on PDMS by *C. albicans/S. aureus* dual-species containing either yeast-form or hyphal-form *C. albicans* cell morphotypes reveals striking differences in biofilm formation on the surface of PDMS. Figure 3 shows biofilm is observed on all PDMS discs, although these differ in degrees of biomass deposition, as illustrated by images representative of biofilm formation in Figure 3 (A) and the quantification of total biofilm coverage within the field of view in Figure 3 (B).

**Figure 3:**
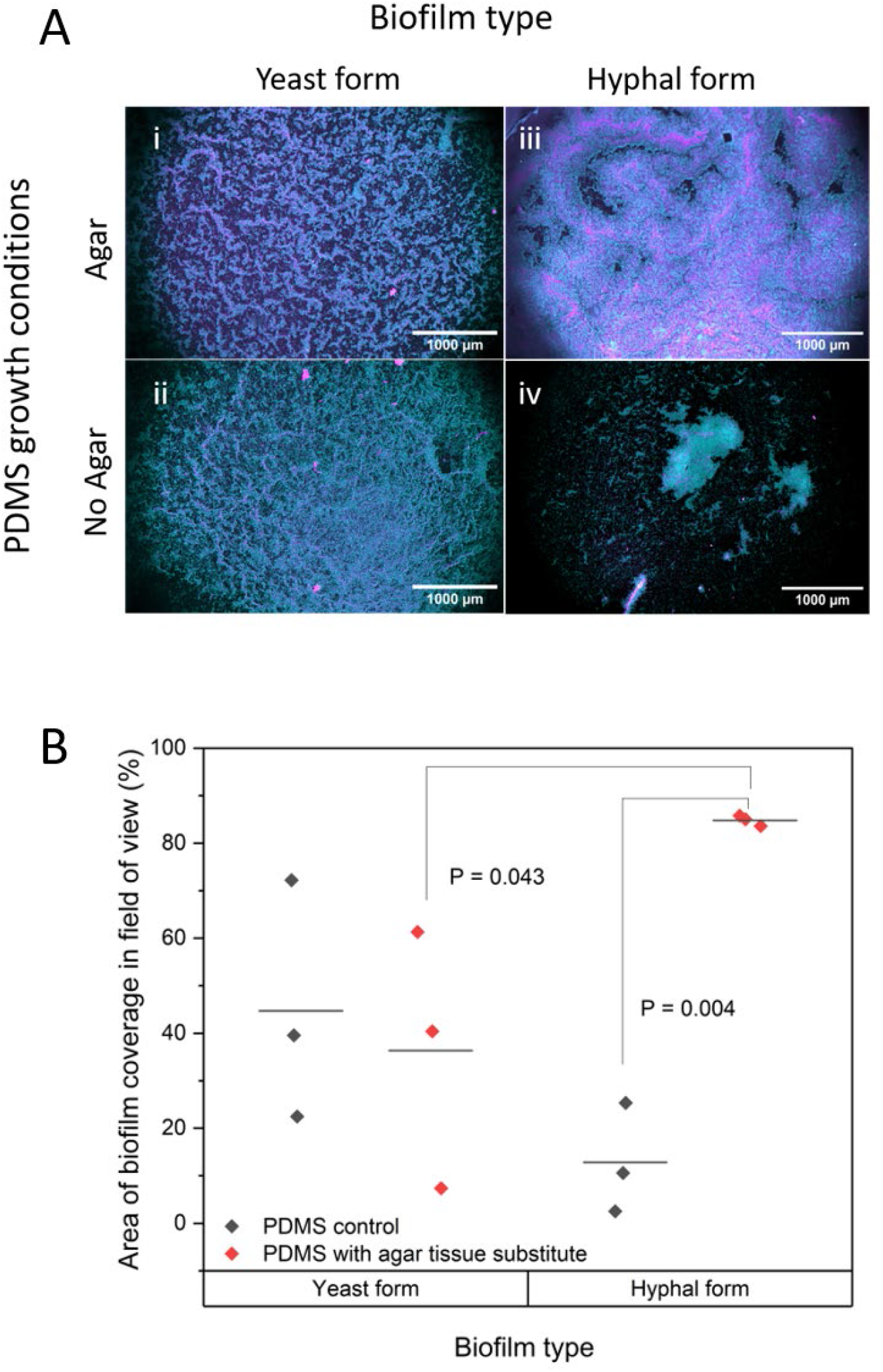
Analysis of biofilm formation on PDMS discs by widefield epifluorescence mesoscopy. A: images of *C. albicans* pACT1-1 GFP (cyan) and *S. aureus* N315 mCherry (magenta) dual species biofilms formed by yeast-form and hyphal-form dual species co-cultures. Panels **(i)** and **(iii)** are images of biofilms grown on PDMS in the presence of agar, and **(ii)** and **(iv)** are images of biofilms grown on PDMS in the absence of agar. Dual-species biofilms grown from predominantly yeast-form *C. albicans* are shown in panels **(i)** and **(ii)**, and biofilms grown from predominantly hyphal-form *C. albicans* containing cultures are shown in panels **(iii)** and **(iv)**. Maximum intensity z projections, scale bars 1000 µm. **B: Quantification of biofilm deposition as a percentage of total surface coverage within the field of view**. Measurement of total biofilm coverage on PDMS discs by C. albicans pACT1-1 GFP and S. aureus N315 mCherry co-cultures. Composite images of both excitation wavelengths were used to quantify total biomass on PDMS as a percentage of the total field of view, under each condition shown. **Black data points:** Biofilms grown on PDMS without agar. **Red data points:** Biofilms grown on PDMS with agar. Statistical analysis by unpaired T-test.

After 48 hours, co-cultures of yeast-form *C. albicans* and *S. aureus* develop similar biofilms on PDMS in both the presence and absence of the agar mimic, as shown in Figure 3A (i) and (ii), respectively, whereas co-cultures of hyphal-form *C. albicans* and *S. aureus* develop biofilms with significantly greater biomass on PDMS mounted in the agar tissue substitute [Figure 3A (iii)] than on PDMS without the agar tissue substitute [Figure 3A (iv)]. These visual observations are confirmed by quantification in Figure 3 (B), which details total biofilm coverage from all three biological replicates of each growth condition. Figure 3 (B) shows biofilms formed by the yeast-form co-culture on PDMS discs in both the presence and absence of the agar tissue substitute do indeed have similar surface coverage, with an average surface percentage of 36% ±19.3% and 45% ±18.3% on agar-mounted and control PDMS discs, respectively. With regards to the hyphal-form co-culture, the marked difference between PDMS discs in the presence and absence of the agar tissue substitute is profound, with an average percentage coverage of 85% ± 0.8% in the presence of agar, in comparison to an average percentage surface coverage of 12% ± 8.3% on the PDMS control discs.

### Deposition and biofilm matrix accumulation by the hyphal-form C. albicans/S. aureus co-culture differ on the silicone elastomer colonisation model

To assess biomass formation by each dual-species co-culture on the silicone elastomer colonisation model, analysis of cell deposition was undertaken by indirect quantification of fluorescence intensity measurements, and the deposition of biofilm matrix components investigated by crystal violet staining. In terms of biomass of cell deposition, the graph in Figure 4 (A) (ii) shows that on average, hyphal-form co-cultures deposit ten-fold greater cell biomass on PDMS in the presence than in the absence of the agar tissue substitute, with an average fold change of 10.14 ± 2.01. Conversely, in biofilms formed by the yeast-form co-culture there is a decrease of approximately 46% ± 21% in cell biomass deposition on PDMS in the presence of the agar tissue substitute in comparison to its absence. This contradicts the data presented in Figure 3 (B), however there is a significant outlier in the yeast-form co-culture datasets presented in Figure 4 (A), which is most likely influencing these averages. Further replicates would confirm whether this observation is indeed true.

**Figure 4:**
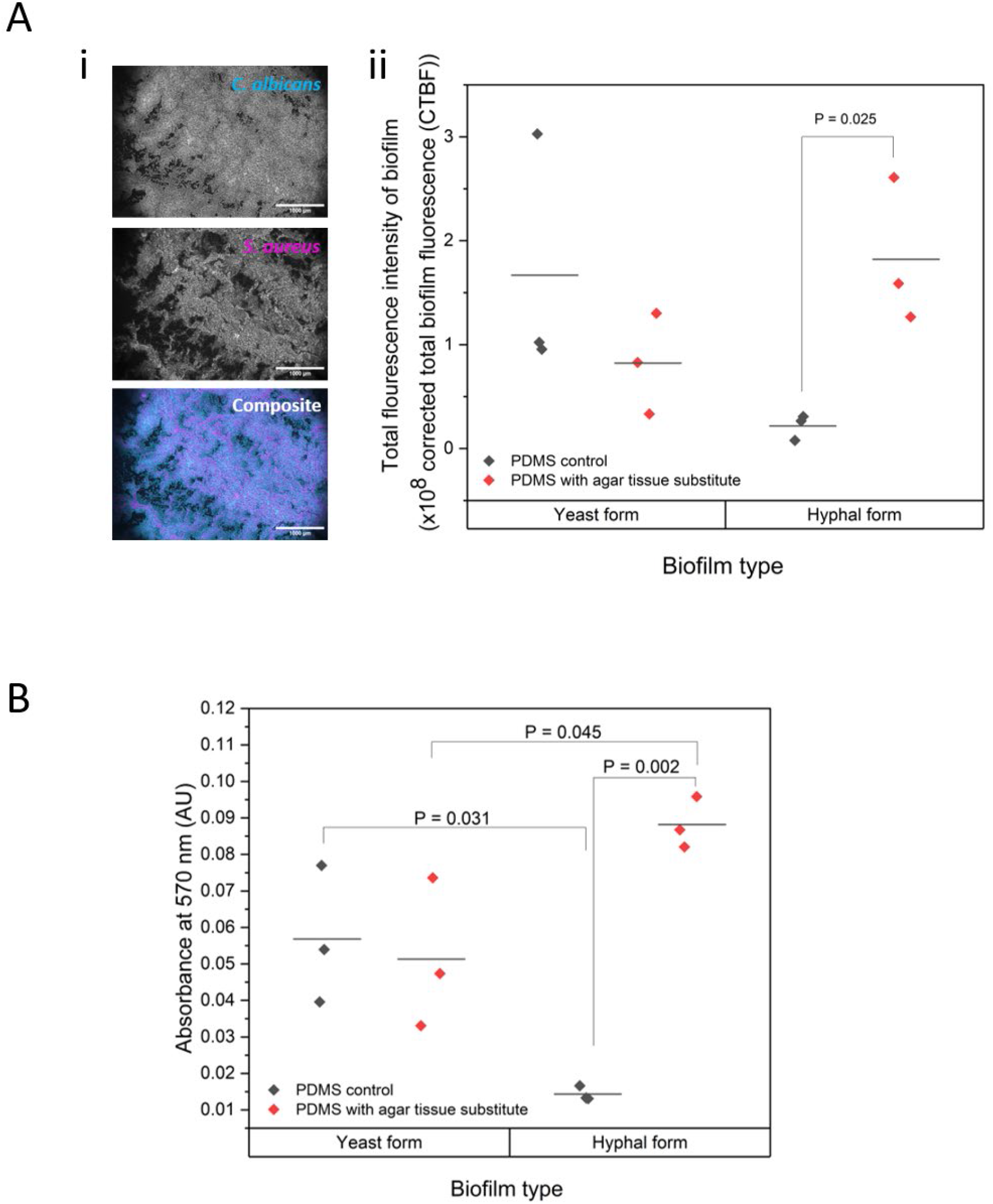
Deposition of biofilm biomass on PDMS. A: Cell deposition indirectly quantified by fluorescence intensity. **(i)** Representative grayscale images of 595 nm channel (*C. albicans*, top panel) and 635 nm channel (*S. aureus*, middle panel) used to calculate combined fluorescence intensities presented in (ii). Bottom panel displays image of combined channels. Maximum intensity Z projections, scale bars 1000 µm **(ii)** Graph of combined fluorescence intensities of both 525 nm and 635 nm channels for each biofilm. **B: Assessment of matrix accumulation by crystal violet assay**. Biofilms quantified by crystal violet staining and measurement at 570 nm. **Black data points:** Biofilms grown on PDMS without agar. **Red data points:** Biofilms grown on PDMS with agar. Statistical analysis by unpaired T-test.

With regard to deposition of biofilm matrix, crystal violet data presented in Figure 4 (B) indicates biofilm matrix deposition on PDMS by the yeast-form *C. albicans/S. aureus* dual-species co-cultures is least influenced by the presence or absence of an agar tissue substitute. Both yeast-form co-culture biofilms average similar absorption values at 570 nm, with a percentage difference of 19.8% ± 12.6%. Biofilm matrix deposition by the hyphal-form *C. albicans/S. aureus* co-culture, however, is heavily influenced by the presence of an agar tissue substitute, averaging a five-fold increase of 525.4% ± 88% in matrix deposition on PDMS discs mounted in agar than those discs without an agar tissue substitute. These hyphal-form dual-species biofilms average absorbances of 0.088 and 0.014 AU, respectively. In terms of the impact of cell morphotype on biofilm matrix deposition, the hyphal-form *C. albicans/S. aureus* co-culture also produced the greatest deposition of matrix on PDMS mounted on the agar tissue substitute, with an 89.5% ± 47.8% increase in deposition in comparison to biofilms formed on the agar-mounted PDMS by the yeast-form *C. albicans/S. aureus* co-culture (average absorbances of 0.088 vs 0.051 AU). Interestingly, both the greatest and the least degree of biofilm matrix and cell deposition on PDMS was produced by the hyphal-form *C. albicans/S. aureus* co-cultures, indicating the critical function of the *C. albicans* hyphal-form cell morphotype in surface colonisation by the *C. albicans/S. aureus* co-culture.

## Discussion

The interplay between *C. albicans* and *S. aureus* in both infection and *in vitro* studies is well documented in the literature (21–25,34,36,37,41,44) however there is a significant lack in our understanding of their synergy in device-associated HAIs. Considering the criticality of HAIs in the global infection burden and their contribution to the emergence of antimicrobial resistance, we aimed to address aspects of that knowledge gap through the development of a simple and inexpensive *in vitro* silicone elastomer colonisation model and applied the system to examine *C. albicans* and *S. aureus* dual-species biofilm formation on a clinically relevant surface. Using previously established methodology for the creation of dual-species biofilms containing either predominantly yeast-form *C. albicans* and *S. aureus*, or hyphal-form *C. albicans* and *S. aureus* (34), this study examined the capacity of these co-cultures to form biofilm on PDMS in the presence or absence of agar acting as a tissue substitute. While agar is neither representative of tissue in terms of rheological properties nor nutrient composition, it is a malleable surface which provides nutritional support. Alternative mounting materials such as hydrogels (45) would provide an environment more similar to that of an indwelling medical device than agar, however, synthesis of these materials is expensive and requires involved preparation-which may be a barrier to those who lack such resources. By creating a simple silicone elastomer colonisation model with common laboratory consumables, this work provides an easily accessible method to explore the surface colonisation of materials used in indwelling medical devices.

Although the hyphal form of *C. albicans* is associated with colonisation of tissues and indwelling medical devices (27,37,38), results from this study revealed yeast-form *C. albicans*/*S. aureus* biofilms readily formed biofilms not only on the PDMS disc of the silicone elastomer colonisation model (SIMCOL), but also on the PDMS control. These yeast-form *C. albicans*/*S. aureus* dual-species biofilms were almost of equivalence when measured by both crystal violet staining and percentage total surface coverage, indicating a tissue substitute-independent attachment capacity. *Candida albicans* surface attachment is mediated in part by the agglutinin-like sequence (ALS) protein family (46), with the hyphal-expressed Als3p playing a key role in both the adhesion of *C. albicans* to surfaces (47) and *S. aureus* interactions (48). This observed ability to form dual-species biofilm in both the presence and absence of an agar tissue substitute could suggest a possible role for other non-Als3p ALS family proteins or other adhesins in surface attachment, since a variety of adhesin molecules are expressed during adherence (46,49–51) Furthermore, contributions from *S. aureus* binding capabilities may also promote formation of the dual-species yeast-form biofilm on PDMS, as a variety of interactions are known to facilitate *S. aureus* surface binding (52),(53). However, the interaction of yeast-form *C. albicans/S. aureus* surface attachments would require further investigation with single species cultures and mutants compromised in protein families associated with adhesion, both of which are out of scope of this study. Surprisingly, the hyphal-form *C. albicans/S. aureus* dual-species co-culture produced very little biofilm on PDMS in the absence of an agar tissue substitute, a stark contrast to the biofilm formed on PDMS in the presence of the agar which yielded the greatest deposition of biofilm of any condition. This observation is unexpected as the hyphal form of *C. albicans* has been shown to attach to PDMS surfaces (27). However, this finding supports the role of Als3p as the key interacting protein with both surfaces and with *S. aureus*, as interactions of *S. aureus* with *C. albicans* Als3p in suspension could competitively inhibit Als3p-mediated surface adhesion resulting in minimal biofilm formation. Again, mutation studies with *als3Δ/als3Δ C. albicans* mutants in hyphal-form dual-species biofilms would shed further light on this finding.

In terms of wider applications of the SIMCOL model, the system has two potential uses. Firstly, it could be employed to explore the behaviour of other clinically relevant microbial communities associated with medical device HAIs, test treatment regimens and systematically probe biofilm attachment and synthesis in response to systematic environmental changes, allowing investigation of the infectious microenvironment (54). Secondly, the SIMCOL model is of benefit to those creating novel materials and approaches in indwelling medical device development. Current research on novel methodologies against medical device associated infections include the modification of surface topologies (55) alterations to surface hydrophobicity (56) functionalisation of surfaces with biosurfactants (57) and surface doping (58–60). Using our simple SIMCOL silicone elastomer colonisation model, investigators developing such technologies would be able to perform inexpensive scoping experiments for technology development, or preliminary data acquisition prior to expansion to other systems for biocompatibility assessment.

## Conclusion

This study presents the first instance of a simple and inexpensive optically tractable novel silicone elastomer colonisation model and demonstrates its application in the assessment of biofilm formation by *C. albicans and S. aureus* on PDMS, a commonly used silicone elastomer material for medical devices. The observed variations in biofilm formation across different conditions underscore the complexity of microbial interactions and suggest avenues for further investigation, particularly into adhesion mechanisms and protein contributions. Beyond the findings presented here, the SIMCOL model offers a valuable tool for exploring microbial dynamics and testing novel antimicrobial strategies, positioning it as a resource for advancing both scientific understanding and technological innovation in the fight against HAIs associated with indwelling medical device infections.

## Acronyms

HAI: Healthcare-associated infections
AMR: antimicrobial resistance
PDMS: Polydimethylsiloxane
LB: Luria Broth
GFP: Green Fluorescent Protein

## Acknowledgements

We would like to thank Professor Ross Fitzgerald (University of Edinburgh) and Dr Fiona Sargison (University of Oxford) for the gift of *S. aureus* strains, and Professor Neil Gow and Professor Alistair Brown (both University of Exeter) for the gift of *C. albicans* strains.

